# Host-specific effects of a generalist parasite of mosquitoes

**DOI:** 10.1101/2024.02.09.579598

**Authors:** Tiago G. Zeferino, Jacob C. Koella

**Affiliations:** Institute of Biology, University of Neuchâtel, Rue Emilie-Argand 11, 2000 Neuchâtel, Switzerland

## Abstract

Microsporidians are obligate parasites of many animals, including mosquitoes. Some microsporidians have been proposed as potential agents for the biological control of mosquitoes and the diseases they transmit due to their detrimental impact on larval survival and adult lifespan. To get a more complete picture of their potential use as agents of biological control, we measured the impact of *Vavraia culicis* on several life-history traits of *Aedes aegypti* and *Anopheles gambiae*. We measured the infection dynamics and clearance rate for the two species, and we assessed sexual dimorphism in infection dynamics within each species. Our results show differences in infection dynamics, with *Ae. aegypti* life-history traits being more affected during its aquatic stage and exhibiting higher clearance of the infection as adults. In contrast, *An. gambiae* was unable to clear the infection. Additionally, we found evidence of sexual dimorphism in parasite infection in *An. gambiae*, with males having a higher average parasite load. These findings shed light and improve our knowledge of the infection dynamics of *V. culicis*, a microsporidian parasite previously recognized as a potential control agent of malaria.

## Introduction

Microsporidians are obligate parasites ^1^ of many animals, particularly insects and fish ^2^, including humans ^3^. Almost half of the species infecting insects use dipterans as hosts ^4^ and more than 150 species have been found in 14 genera of mosquitoes ^5–12^. While their life cycles vary from simple cycles in a single host to complex cycles involving several species ^11^, they are all transmitted to larval mosquitoes and kill some larvae, letting others survive to cause, in adults, lower fertility, shorter lifespan and loss of vigour ^13,14^.

A well-studied example is *V. culicis*, an intracellular parasite which infects several genera of mosquitoes ^15^. Often studied in *Aedes* (the vectors of arboviruses like dengue or Zika) and *Anopheles* (the vector of malaria), it infects larvae when they ingest the spores along with their food. After several rounds of replication, that take about 10 days of development, the parasite starts to produce new infectious spores that spread to gut and fat body cells. In nature, transmission happens through the release of spores in the aquatic environment from infected larvae or adults, from mother to offspring through spores adhering to the surface of the eggs, or, less commonly, from spores released via faeces. However, since the parasite is mildly pathogenic and generally kills few larvae, larva-to-larva transmission is probably rare ^6^.

The accumulation of spores is correlated with greater host mortality ^16^. Because in addition to killing mosquitoes *V. culicis* impedes the development of malaria in mosquitoes ^15^, it (and other microsporidians like microsporidia MB ^12,17,18^) has long been suggested as a way to slow the transmission of malaria ^19^. This is becoming particularly relevant in the face of increasing resistance against chemical insecticides, for *V. culicis* also restores, at least partially, the sensitivity to insecticides of genetically resistant mosquitoes ^20^.

An important feature of using *V. culicis* to control malaria is that this microsporidian in *Anopheles* mosquitoes has little impact on the survival of young adults, but kills mostly old mosquitoes ^21^, in particular when they are infected by malaria ^22^. It has therefore been suggested that there is only little selection pressure for resistance and thus that it can be used in an evolutionarily more sustainable way than chemical insecticides.

The ability of *V. culicis* to control *Aedes*-borne viruses in an evolutionarily sustainable way is less clear. Indeed, since the impact of *V. culicis* on *Aedes* ^9,10,23^ is greater than on *Anopheles* and, in particular, since infection increases the rate of mortality in all age classes ^24^, selection pressure for resistance to *Vavraia* is expected to be higher for *Aedes* than that for *Anopheles*.

There are, however, many gaps in our knowledge about the biology of *V. culicis*, both in *Anopheles* and in other mosquitoes, in particular since traits other than mortality rates have only rarely been investigated. To get a complete picture of the potential use of *V. culicis* as an agent of biological control ^25^, we compared the impact of the parasite on several life-history traits of *Ae. aegypti* and *An. gambiae*, we measured the infection dynamics and clearance rate for the two species, and we assessed sexual dimorphism in infection dynamics within each species.

## Results

### 1. Larvae and pupae mortality

*Ae. aegypti* and *An. gambiae* larvae were reared individually in 12-well culture plates, each containing 3 ml of deionized water per well. Two-day-old larvae were either exposed to *V. culicis* or left unexposed. This experimental design allowed us to disentangle larvae and pupae mortality, which were assessed every 24 hours.

Larval mortality was higher in *An. gambiae* than *Ae. aegypti* (0.36 vs 0.02, χ^2^ = 523.82, df = 1, p < 0.001, **Fig. 1a**) and exposure to *V. culicis* increased larval mortality (χ^2^ = 112.25, df = 1, p < 0.001, **Fig. 1a**) by a factor of about 2.3 in *An. gambiae* and 3.7 in *Ae. aegypti*.

**Figure 1.**
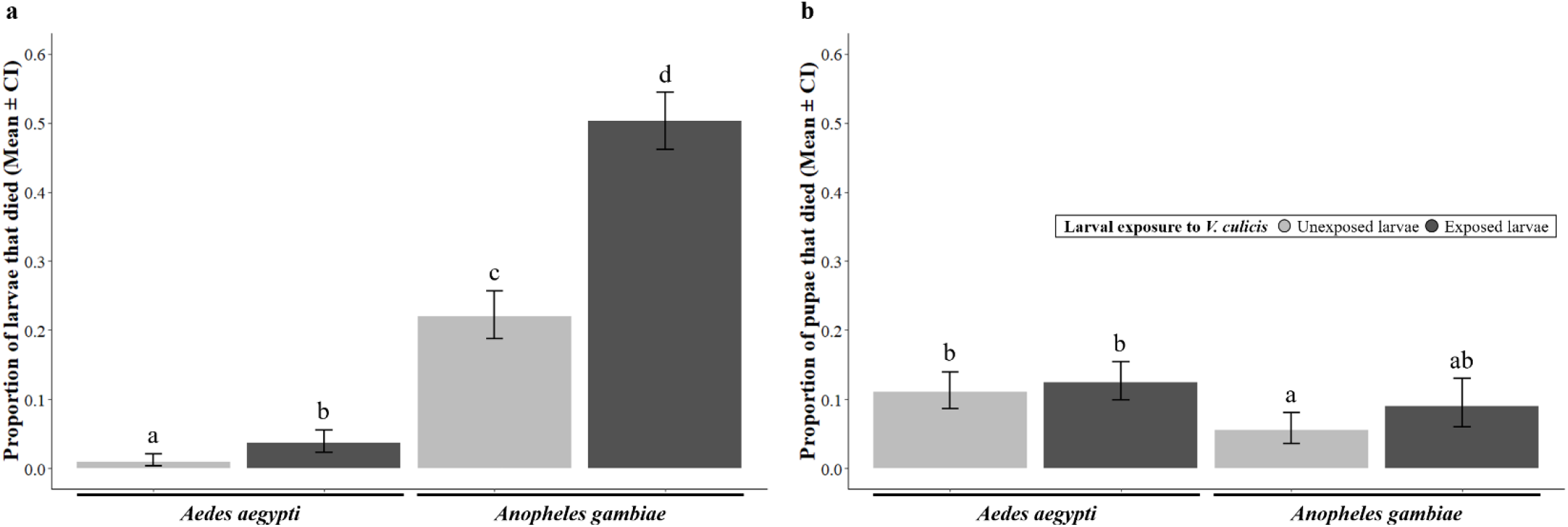
Larvae and pupae mortality. The proportion of *Ae. aegypti* and *An. gambiae*, exposed or unexposed to *V. culicis*, that died **(a)** as larvae or **(b)** as pupae throughout the aquatic stage. The sample sizes were 600, 590, 576 and 578 for larvae, and 594, 568, 449 and 287 for pupae. Error bars show the 95% confidence intervals of the means. Letters indicate statistically significant differences from multiple comparisons.

Pupal mortality was higher in *Ae. aegypti* than *An. gambiae* (0.12 vs 0.07, χ^2^ = 11.34, df = 1, p < 0.001, **Fig. 1b**), but was not affected by exposure to *V. culicis* (χ^2^ = 2.47, df = 1, p = 0.115, **Fig. 1b**). For further details on the statistical analysis see **Supplementary Table S1**.

### 2. Age at pupation, wing length and sex ratio

Age at pupation was assessed every 24 hours, and pupae were transferred individually to falcon tubes. Upon emergence, we measured wing length and determined the sex of each mosquito.

*An. gambiae* took about one day longer to pupate than *Ae. aegypti* (8.51 days vs 7.48 days, F_1,1702_ = 298.32, df = 1, p < 0.001), and females took longer than males (8.17 days vs 7.82 days, F_1,1702_ = 58.88, df = 1, p < 0.001). However, the effect of sex was only evident for *Ae. aegypti* (interaction sex * species: F_1,1702_ = 6.25, df = 1, p = 0.012, **Fig. 2a**). Exposed individuals pupated, on average, half a day later than unexposed ones (8.26 days vs 7.72 days, F_1,1702_ = 147.90, df = 1, p < 0.001), but the effect of exposure was only significant in *Ae. aegypti* (interaction species * exposure: F_1,1702_ = 36.56, df = 1, p < 0.001, **Fig. 2a**).

**Figure 2.**
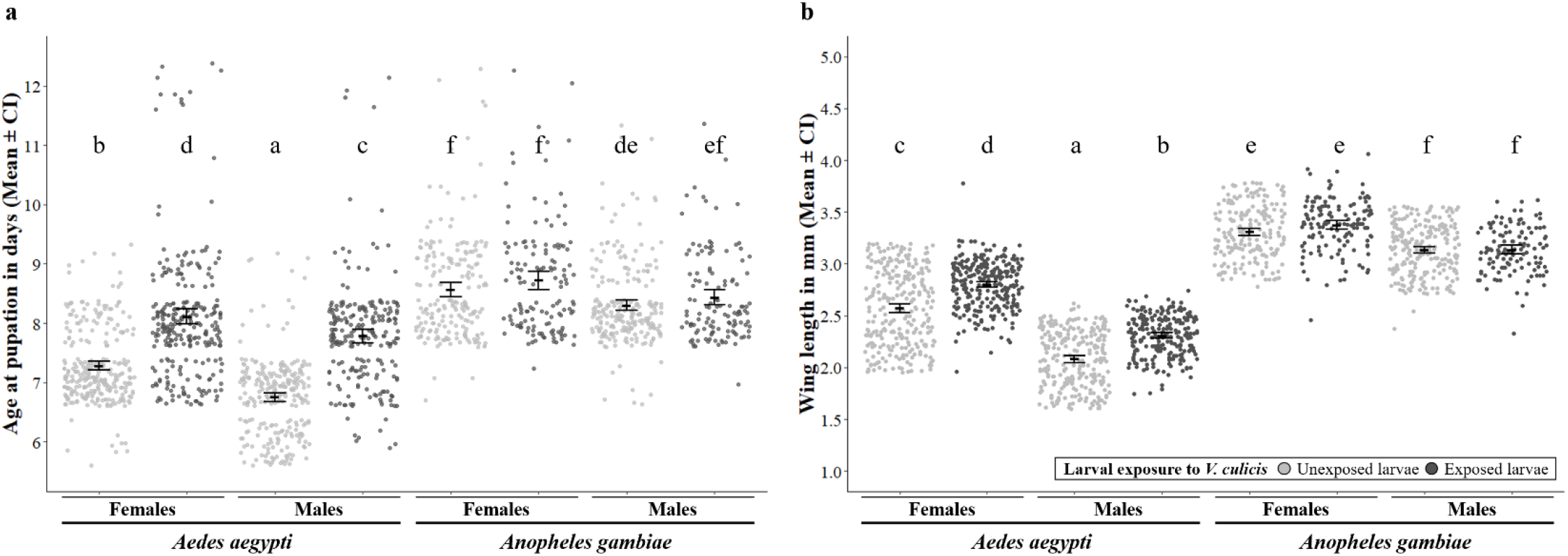
Age at pupation and wing length. **(a)** Age at pupation in days and **(b)** wing length in mm of males and females, exposed or not to *V. culicis*, for *Ae. aegypti* and *An. gambiae*. The sample sizes were 275, 269, 253, 228, 195, 137, 229, and 124. Error bars show the 95% confidence intervals of the means. Letters indicate statistically significant differences from multiple comparisons.

*An. gambiae* had longer wings than *Ae. aegypti* (3.24 mm vs 2.44 mm, F_1,1702_ = 880.08, df = 1, p < 0.001), and females had longer wings than males (3.01 mm vs 2.67 mm, F_1,1702_ = 448.23, df = 1, p < 0.001). However, the effect of sex was stronger for *Ae. aegypti* (22% longer wings vs 6% longer wings, interaction sex * species: F_1,1702_ = 82.88, df = 1, p < 0.001, **Fig. 2b**). Exposed individuals had longer wings than unexposed ones (2.90 mm vs 2.77 mm, F_1,1702_ = 100.46, df = 1, p < 0.001), but the effect of exposure was only significant in *Ae. aegypti* (interaction species * exposure: F_1,1702_ = 18.62, df = 1, p < 0.001, **Fig. 2b**). For further details on the statistical analysis see **Supplementary Table S2**.

The proportion of females among the adults was higher for *Ae. aegypti* than for *An. gambiae* (0.53 vs 0.48, χ^2^ = 14.61, df = 1, p < 0.001), and higher for exposed individuals than for unexposed ones (0.53 vs 0.49, χ^2^ = 28.63, df = 1, p < 0.001). In addition, the proportion of females also increased with later pupation (from 0.08 at day 6 to 0.81 at day 12, χ^2^ = 97.00, df = 1, p < 0.001), and this effect was influenced by species and exposure (interaction age at pupation * species * exposure: χ^2^ = 15.10, df = 1, p < 0.001, **Fig. 3**). Thus, while in *An. gambiae* the sex ratio was only influenced by age at pupation, in *Ae. aegypti* the shift from producing more males to producing more females was delayed by 24h in exposed mosquitoes. For further details on the statistical analysis see **Supplementary Table S3**.

**Figure 3.**
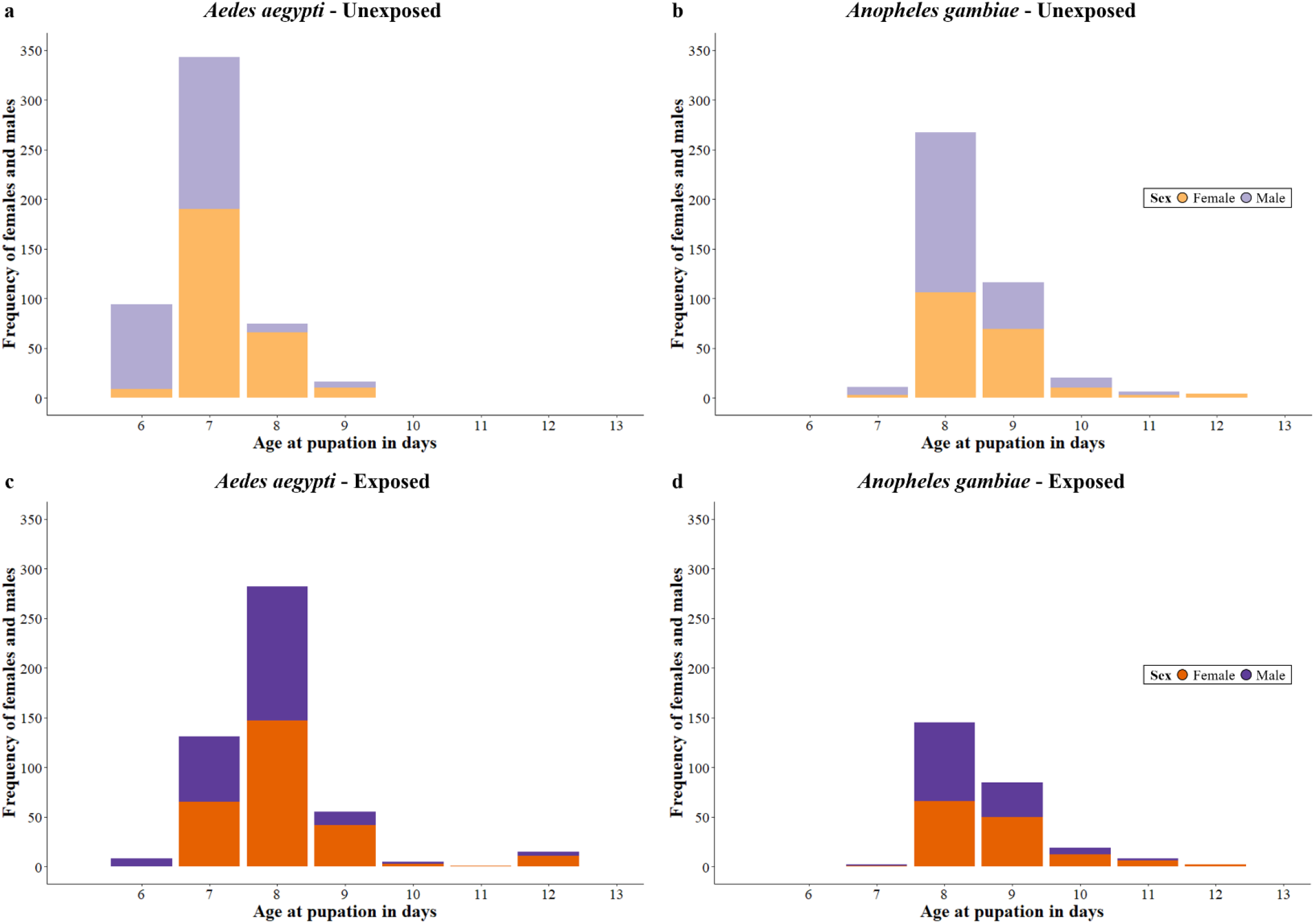
Sex ratio. Frequency of females and males of **(ac)** *Ae. aegypti* and **(bd)** *An. gambiae*, exposed or not to *V. culicis*, for each age at pupation. The sample sizes were 528, 497, 424 and 261 respectively.

### 3. Presence of spores and spore density

We assessed infection dynamics by sampling 60 mosquitoes (30 males and 30 females) every other day, starting on the day of emergence. Mosquitoes were killed, and the number of spores was counted with a haemocytometer containing 0.1 μl of the sample.

In *Ae. aegypti*, the proportion of individuals harbouring spores increased for the first 10 days and then decreased with the age of the mosquitoes (χ^2^ = 3.99, df = 1, p = 0.046, **Fig. 4a**), showing an ability to clear the infection independently of sex (χ^2^ = 1.28, df = 1, p = 0.258). Spore density (on mosquitoes harbouring spores) increased with the age of the mosquitoes (χ^2^ = 19.68, df = 1, p < 0.001) up to a plateau of about 10^4^ to 10^5^ spores per mosquito. While spore density was similar for males and females (χ^2^ = 1.07, df = 1, p = 0.300), sex influenced the increase with age (interaction age * sex: χ^2^ = 6.07, df = 1, p = 0.014, **Fig. 5a**), so that by the end of their lives, males had a slightly higher parasite load than females.

**Figure 4.**
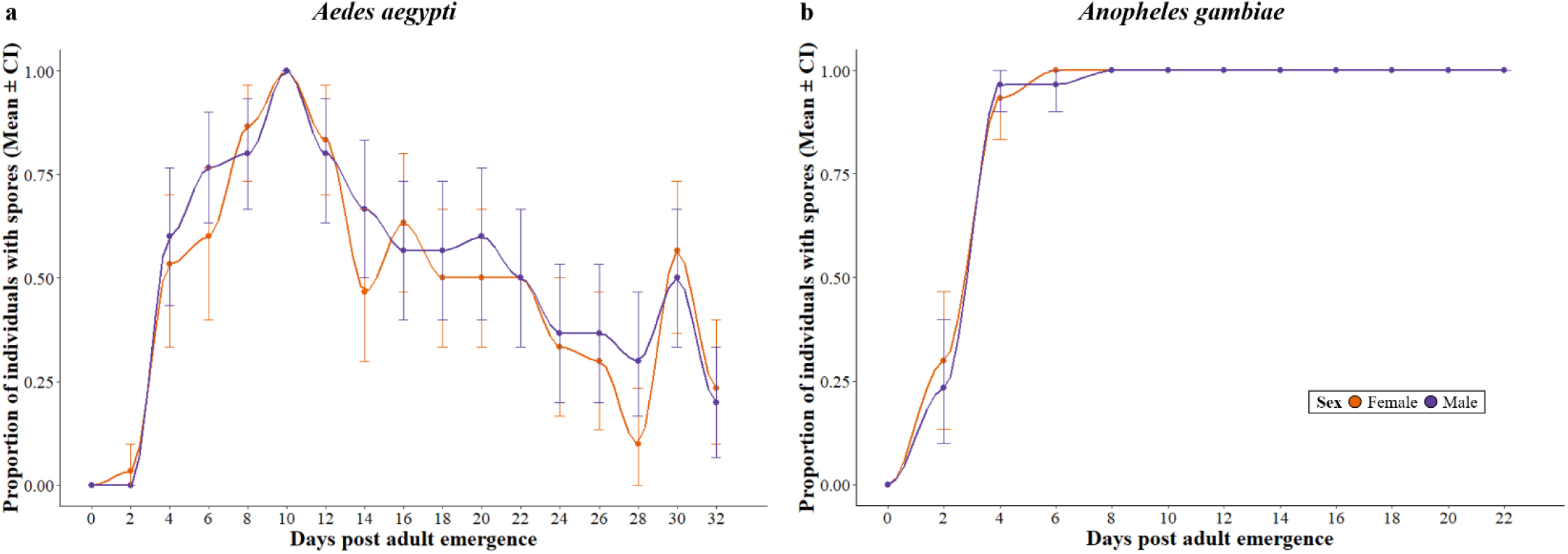
Presence of spores. The proportion of alive individuals that had detectable spores for each sex for **(a)** *Ae. aegypti* and **(b)** *An. gambiae*. The sample size was 30 individuals per time point per sex. Error bars show the 95% confidence intervals of the means. A smooth line, calculated with the “loess” method of ggplot2, was drawn through the means for ease of visualization.

**Figure 5.**
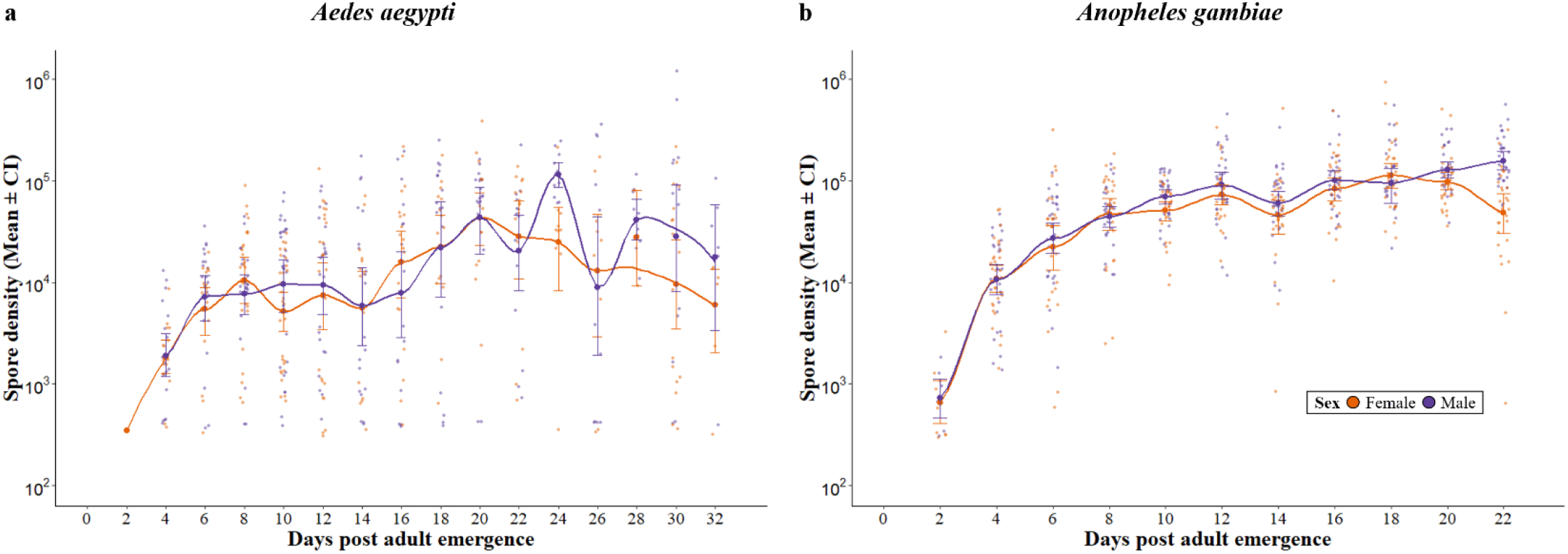
Spore density. Spore density for **(a)** *Ae. aegypti* and **(b)** *An. gambiae*. Mosquitoes without any spores are not shown. Error bars show the 95% confidence intervals of the means. A smooth line, calculated with the “loess” method of ggplot2, was drawn through the means for ease of visualization.

In *An. gambiae*, the proportion of individuals harbouring spores increased with the age of the mosquitoes (χ^2^ = 501.28, df = 1, p < 0.001, **Fig. 4b**) until it reached 100% at day 8, and then remained constant. This pattern was the same for males and females (χ^2^ = 0.25, df = 1, p = 0.618). Spore density increased with the age of the mosquitoes (χ^2^ = 165.84, df = 1, p < 0.001) up to about 10^5^ spores, and males harboured more spores than females (χ^2^ = 7.22, df = 1, p = 0.007). There was no interaction between age and sex (χ^2^ = 3.44, df = 1, p = 0.064, **Fig. 5b**). For further details on the statistical analysis see **Supplementary Table S4**.

## Discussion

Although we had alternated the transmission of *V. culicis* through its two mosquito hosts for several years before this experiment, it affected *Ae. aegypti* more than *An. gambiae* but could be cleared only by the former.

Juvenile mortality was considerably higher for infected than for uninfected mosquitoes. While this result is not novel ^26^, it is noteworthy in several respects. First, in line with our general observation, the impact of infection was about 60% greater for *Ae. aegypti* than for *An. gambiae*. Second, we distinguished between larval and pupal mortality (**Fig. 1**). Indeed, most of the parasite-induced pre-adult mortality occurred in larvae, while pupal mortality was not affected by infection, in contrast to suggestions that *V. culicis* is a pupicidal parasite ^24,27^. These results are consistent with the idea that horizontal transmission among larvae is the main mode of transmission of *V. culicis*, even in situations where (like in this experiment) larvae are well-fed. This contrasts the suggestion that the virulence of *V. culicis* is low when mosquitoes are well-fed ^28–30^. This difference may be because we reared mosquitoes individually, thus avoiding competition among individuals, which can increase mortality ^31^ and could confound the impact of infection.

Time to pupation and wing length were also changed by infection, and again this change was apparent only in *Ae. aegypti* (**Fig. 2**). Infected mosquitoes pupated later than uninfected mosquitoes, corroborating several studies on the impact of microsporidians on mosquitoes ^15,24,32^, although in at least one study infected mosquitoes pupated earlier than uninfected ones ^28^. This result is what is expected for hosts that change their life history to weaken the impact of larval infection on reproductive success. More unexpected, both in terms of theory ^33,34^ and in comparison with other studies ^15^, is that infected adults were larger than uninfected ones. The mechanism underlying these differences among studies and species are not known.

In infected mosquitoes, the larval development can be expected to change according to the host’s or the parasite’s interest. On one hand, parasites could manipulate their host’s development to increase their transmission in at least two ways. They could increase horizontal transmission by delaying pupation, thus having more time to produce infectious spores. If infected larvae grow at the same rate as uninfected ones, manipulation for later pupation would bring with it larger adults. In this experiment, we observed both of these changes for *Ae. aegypti*. While this corroborates most studies on the impact of microsporidians on mosquitoes concerning age pupation ^15,24,32^, infected individuals are smaller than uninfected ones in most studies ^15^, as expected if infection slows growth. In addition, since males cannot transmit the parasite vertically, the parasite could also selectively kill male larvae to take advantage of their role in horizontal transmission. This would bias the sex ratio of adults towards females, as observed here for *Ae. aegypti* (**Fig. 3**). On the other hand, the host should evolve to decrease the impact of the parasite on its reproductive success. This would favour earlier pupation at a smaller size of infected individuals ^33,34^, as observed for microsporidian-infected *Culex-*mosquitoes ^28^. Finally, since females take longer to develop than males, the parasite has more time to kill females than males during larval development. This constraint would therefore lead to male-biased adults, as observed for *Edhazardia*- infected *Ae. aegypti* ^35^. Overall, thus, the impact of the parasite on the host’s development is determined by a balance between the host’s interests, the parasite’s interests and developmental constraints. While in our experiment *V. culicis* appeared to overwhelm the interests of *Ae. aegypti*, in many cases the host’s interests dominate the interaction.

The proportion of individuals with detectable spores and the density of the spores were the traits most influenced by the species (**Fig. 4** and **5**). The proportion of individuals with detectable spores increased with their age for *An. gambiae*, while for *Ae. aegypti*, it reached a peak after 10 days and then decreased with age. This suggests that, *Ae. aegypti* (but not *An. gambiae*) was able to clear the infection, in particular as they became older. The ability of *Ae. aegypti* to clear the parasite was reflected in the low density of spores in many individuals. This led to greater variance and a lower average of the density of spores in *Ae. aegypti* than in *An. gambiae*, although the maximal densities in the two species were similar. We also found sexual dimorphism in infection dynamics, with males harbouring on average more spores than females, as is generally observed ^36–42^.

Our results provide a more comprehensive understanding of the potential use of *V. culicis* as a biological control agent ^25^. On one hand, it strengthens the argument of using *V. culicis* to slow ^19^ or impede ^15^ the transmission of malaria. A particular advantage is that, in our study, *An. gambiae* did not clear *V. culicis*, allowing it to impact malaria transmission throughout the mosquito’s life. On the other hand, the potential use for malaria control cannot be extrapolated to diseases such as dengue, which are transmitted by *Ae. aegypti*. Not only is the parasite more virulent in *Ae. aegypti* than in *An. gambiae* ^24^, making it likely that resistance evolves rapidly, but it is also cleared by *Ae. aegypti*.

Our study also highlights the difficulty in predicting how a generalist parasite behaves in its different species of hosts. *V. culicis* is often considered to be a generalist, for it can complete its life cycle in many species of mosquitoes. Moreover, in our laboratory, it is selected to be a generalist, for it is forced to alternate its generations in *Ae. aegypti* and in *An. gambiae*. Nevertheless, its impact on the two mosquitoes differs strongly, with more damage but also more resistance in *Ae. aegypti* than in *An. gambiae*. Yet, it is difficult to know whether it is more specialized on one than on the other, without further experiments quantifying the total transmission through the two mosquitoes. The problem of knowing where on the spectrum of generalist to specialist makes it difficult to interpret studies making predictions about generalist and specialist parasites in epidemiological and ecological studies ^43,44^.

Finally, laboratory tests have assessed the host ranges of many microsporidia species, primarily to ensure the safety of these pathogens for biological control programs ^45,46^. However, in natural settings, host availability may be seasonal, and host abundance can vary throughout the year. Multiple or alternate hosts can act as reservoirs for these microsporidia, enabling their persistence during periods of low host population density ^47^. Considering the distinct ecological preferences and limited distribution overlap of *Ae. aegypti* and *An. gambiae*, it would be intriguing to assess *V. culicis*’s potential to adapt and specialize by evolving it over several generations exclusively within each host. Rapid adaptation of *V. culicis* to its host could restore its disease control potential for diseases transmitted by *Ae. aegypti*.

## Materials and Methods

### Experimental system

We used the UGAL strain of the mosquito *Ae. aegypti* (obtained from Patrick Guérin, University of Neuchâtel) and the Kisumu strain of *An. gambiae s*.*s* ^48^. We maintained the two colonies at a density of about 600 individuals per cage and standard lab conditions (26 ± 1ºC, 70 ± 5% relative humidity and 12 h light/dark) for several years before the experiments.

*V. culicis floridensis* was provided by J.J Becnel (USDA, Gainesville, FL, USA). It is an obligatory, intracellular parasite. Mosquitoes are exposed to *V. culicis* during the larval aquatic stage where they ingest the spores along with their food. After several rounds of replication, the parasite produces new infectious spores that spread to gut and fat body cells. In nature, transmission happens through the release of spores in the aquatic environment from infected larvae or adults, from mother to offspring through spores adhering to the surface of the eggs, or, less commonly, from spores released via faeces. However, since the parasite is mildly pathogenic and generally kills few larvae, larva-to-larva transmission is probably rare ^6^.

The parasite was first found in *Ae. albopictus*, and was later found to parasitize several mosquito genera, including *Aedes, Culex, Anopheles, Culiseta, Ochlerotatus* and *Orthopodomyia* ^49,50^. To ensure that it remains a generalist parasite, we maintained the colony by alternately infecting *Ae. aegypti* and *An. gambiae*. We started by infecting two-day-old larvae in groups of 50 individuals. Upon pupation, the individuals were moved to cages, where they emerged and were fed with cotton balls soaked in 6% sucrose solution, which were replaced daily. Dead individuals were collected every other day, placed into 2 ml Eppendorf tubes containing 1 ml of deionised water and a stainless-steel bead (∅ 5 mm), and stored at 4º C. These individuals (around 15 per tube) were later homogenised with a Qiagen TissueLyser LT at a frequency of 30 hz for two minutes, and used to infect new larvae.

### Mosquito rearing and maintenance

Freshly hatched (0-3h old) *Ae. aegypti* and *An. gambiae* larvae were put individually into 12-well culture plates containing 3 ml of deionized water per well. The larvae were fed daily with Tetramin Baby® fish food according to their age (*Ae. aegypti*: 0.06, 0.08, 0.16, 0.32, 0.64 and 0.32 mg/larva respectively on days 0, 1, 2, 3, 4 and 5 or older ^51^; *An. gambiae*: 0.04, 0.06, 0.08, 0.16, 0.32 and 0.6 mg/larva respectively on days 0, 1, 2, 3, 4 and 5 or older ^52^).

We exposed two-day-old larvae to 0 or 20,000 (*Ae. aegypti*) or to 10,000 (*An. gambiae*) spores of *V. culicis*. Pupae were moved to individual 50 ml falcon tubes containing about 10 ml of deionized water ^53^, and adults were moved to 150 ml cups (5 cm ∅ x 10 cm) that were covered with a net and contained a 55 mm ∅ petri dish (to keep the mosquitoes from drowning) on the surface of 50 ml deionized water and a 10 x 7 cm rectangular filter paper (to keep the air humid). They were fed with cotton balls soaked with 6% sucrose solution, which were replaced daily.

### Life-history measurements

For each individual, we measured larval and pupal mortality and (for the mosquitoes that emerged) age at pupation, wing length and sex. Larval and pupal mortality and age at pupation were assessed every 24h. The right wing of every mosquito was dissected and the distance from the axillary incision to the tip of the wing ^54,55^ was measured using the software ImageJ v1.54 ^56^.

We assessed infection dynamics by killing 60 mosquitoes every other day, starting on the day of emergence, until there were no more living mosquitoes. We put each mosquito into a 2 ml Eppendorf tube containing 0.1 ml of deionised water, added a stainless-steel bead (∅ 5 mm) to each tube and homogenised the samples with a Qiagen TissueLyser LT at a frequency of 30 hz for two minutes, and counted the number of spores in a haemocytometer containing 0.1 μl of the sample under a phase-contrast microscope (400x magnification).

For the infection dynamics, we calculated the density of spores (that is, the spore count normalized for the size of the mosquito) by dividing the number of spores by the wing length (mm). This gives what we call spore density. Note that, since eight (*Aedes*) and ten (*Anopheles*) days after emergence all mosquitoes harboured spores, the absence of spores in older mosquitoes can also be considered to be the rate at which the mosquitoes can clear the parasite.

### Statistical analysis

All analyses were done with the R software ^57^ version 4.3.0, using the packages “DHARMa” ^58^, “car” ^59^, “lme4” ^60^, “emmeans” ^61^ and “multcomp” ^62^. Significance was assessed with the “Anova” function of the “car” package ^59^. We used a type III ANOVA in the case of a significant interaction and a type II ANOVA otherwise. When relevant, we performed post-hoc multiple comparisons with the package “emmeans”, using the default Tukey adjustment.

Larval and pupal mortality were analysed with a generalized linear model with a binomial distribution of errors, where the response variable was the proportion of dead individuals for each developmental stage and the explanatory variables were species and infection status.

Age at pupation and wing length were analysed with a linear model with a Gaussian distribution of errors, where the response variable was time to pupae or wing length and the explanatory variables were species, infection status and sex.

The sex ratio was analysed with a generalized linear model with a binomial distribution of errors, where the response variable was the sex of the mosquito and the explanatory variables were time to pupae and infection status.

The presence of spores was analysed with a generalized linear model with a binomial distribution of errors, where the response variable was the presence or absence of detectable spores and the explanatory variables were the individual’s age and sex. Because we counted the number of spores in a haemocytometer containing 0.1 μl of the sample (i.e. 1/1000 of the total volume), the detection threshold was estimated to be 1000 spores.

Spore density of the mosquitoes with positive spore counts was analysed with a generalized linear model with a quasi-Poisson distribution of errors, where the response variable was spore density and the explanatory variables were the individual’s age and sex.

## Supporting information

Supplementary information for "Host-specific effects of a generalist parasite of mosquitoes" by Tiago G. Zeferino and Jacob C. Koella

## Acknowledgements

We thank Luís M. Silva for his advice and technical support. We also thank two anonymous reviewers for their insightful and constructive comments. TGZ and the project were supported by SNF grant 310030_192786.

## Author contributions

TGZ conceived the overall idea. TGZ and JCK designed the experiments. TGZ collected the data. TGZ analysed the data and wrote the first draft of the manuscript. All authors contributed critically to the drafts.

## Data availability

All data generated or analysed during this study are included as Supplementary Information files.

## Additional Information

The authors declare no competing interests.

## References

1. Weidner, E. & Overstreet, R. Microsporidia have a peculiar outer membrane with exterior cytoplasmic proteins. Acta Parasitol 1–6 (2021).

2. Franzen, C. Microsporidia: a review of 150 years of research. Open Parasitol J 2, 1–34 (2008).

3. Desportes, I. et al. Occurrence of a new microsporidan: Enterocytozoon bieneusi n.g., n. sp., in the enterocytes of a human patient with AIDS. J Protozool 32, 250–254 (1985).

4. Becnel, J. J. & Andreadis, T. G. Microsporidia in insects. Microsporidia: pathogens of opportunity 521–570 (2014).

5. Bukhari, T., Pevsner, R. & Herren, J. K. Microsporidia: a promising vector control tool for residual malaria transmission. Frontiers in Tropical Diseases 3, 957109 (2022).

6. Andreadis, T. G. Microsporidian parasites of mosquitoes. J Am Mosq Control Assoc 23, 3–29 (2007).

7. Fox, R. M. & Weiser, J. A microsporidian parasite of Anopheles gambiae in Liberia. J Parasitol 45, 21–30 (1959).

8. Gajanana, A. et al. Partial suppression of malaria parasites in Aedes aegypti and Anopheles stephensi doubly infected with Nosema algerae and Plasmodium. Indian Journal of Medical Research 70, 417–423 (1979).

9. Hulls, R. H. Studies on microsporida of mosquitoes and their relationship with Plasmodium berghei. Doctoral dissertation, University of London. (1972).

10. Hulls, R. H. The adverse effects of a microsporidan on sporogony and infectivity of Plasmodium berghei. Trans R Soc Trop Med Hyg 65, 421–422 (1971).

11. Andreadis, T. G. & Hall, D. W. Development, ultrastructure, and mode of transmission of Amblyospora sp. (Microspora) in the mosquito. J Protozool 26, 444–452 (1979).

12. Herren, J. K. et al. A microsporidian impairs Plasmodium falciparum transmission in Anopheles arabiensis mosquitoes. Nat Commun 11, 2187 (2020).

13. Brooks, W. M., Becnel, J. J., & Kennedy, G. G. Establishment of Endoreticulatus NG for Pleistophora fidelis (Hostounský & Weiser, 1975)(Microsporida: Pleistophoridae) Based on the Ultrastructure of a Microsporidium in the Colorado Potato Beetle, Leptinotarsa decemlineata (Say)(Coleoptera: Chrysomelidae) 1, 2. The Journal of protozoology, 35(4), 481–488 (1988).

14. Solter, L. F., Becnel, J. J. & Vávra, J. Research methods for entomopathogenic microsporidia and other protists. Manual of techniques in invertebrate pathology 12, 329–371 (2012).

15. Bedhomme, S., Agnew, P., Sidobre, C. & Michalakis, Y. Virulence reaction norms across a food gradient. Proc R Soc Lond B Biol Sci 271, 739–744 (2004).

16. Agnew, P., Berticat, C., Bedhomme, S., Sidobre, C. & Michalakis, Y. Parasitism increases and decreases the costs of insecticide resistance in mosquitoes. Evolution (N Y) 58, 579–586 (2004).

17. Nattoh, G. et al. Horizontal transmission of the symbiont Microsporidia MB in Anopheles arabiensis. Front Microbiol 12, 647183 (2021).

18. Knell, R. J. & Webberley, K. M. Sexually transmitted diseases of insects: distribution, evolution, ecology and host behaviour. Biological Reviews 79, 557–581 (2004).

19. UNICEF. et al. Fourth Meeting of the Scientific Working Group on Biological Control of Insect Vectors of Disease. No. TDR/BCV-SWG (4)/80.3. World Health Organization (1980).

20. Koella, J. C., Saddler, A. & Karacs, T. P. S. Blocking the evolution of insecticide-resistant malaria vectors with a microsporidian. Evol Appl 5, 283–292 (2012).

21. Koella, J. C., Lorenz, L. & Bargielowski, I. Microsporidians as evolution-proof agents of malaria control? Adv Parasitol 68, 315–327 (2009).

22. Bargielowski, I. & Koella, J. C. A possible mechanism for the suppression of Plasmodium berghei development in the mosquito Anopheles gambiae by the microsporidian Vavraia culicis. PLoS One 4, e4676 (2009).

23. Schenker, W., Maier, W. A. & Seitz, H. M. The effects of Nosema algerae on the development of Plasmodium yoelii nigeriensis in Anopheles stephensi. Parasitol Res 78, 56–59 (1992).

24. Michalakis, Y. et al. Virulence and resistance in a mosquito–microsporidium interaction. Evol Appl 1, 49–56 (2008).

25. Lorenz, L. M. & Koella, J. C. The microsporidian parasite Vavraia culicis as a potential late life–acting control agent of malaria. Evol Appl 4, 783–790 (2011).

26. Sy, V. E., Agnew, P., Sidobre, C. & Michalakis, Y. Reduced survival and reproductive success generates selection pressure for the dengue mosquito Aedes aegypti to evolve resistance against infection by the microsporidian parasite Vavraia culicis. Evol Appl 7, 468–479 (2014).

27. Rivero, A., Agnew, P., Bedhomme, S., Sidobre, C. & Michalakis, Y. Resource depletion in Aedes aegypti mosquitoes infected by the microsporidia Vavraia culicis. Parasitology 134, 1355–1362 (2007).

28. Agnew, P., Bedhomme, S., Haussy, C. & Michalakis, Y. Age and size at maturity of the mosquito Culex pipiens infected by the microsporidian parasite Vavraia culicis. Proc R Soc Lond B Biol Sci 266, 947–952 (1999).

29. Reynolds, D. G. Laboratory studies of the microsporidian Plistophora culicis (Weiser) infecting Culex pipiens fatigans Wied. Bull Entomol Res 60, 339–349 (1970).

30. Reynolds, D. G. Experimental introduction of a microsporidian into a wild population of Culex pipiens fatigans Wied. Bull World Health Organ 46, 807 (1972).

31. Hauser, G., Thiévent, K. & Koella, J. C. Consequences of larval competition and exposure to permethrin for the development of the rodent malaria Plasmodium berghei in the mosquito Anopheles gambiae. Parasit Vectors 13, 1–11 (2020).

32. Bedhomme, S., Agnew, P., Sidobre, C. & Michalakis, Y. Sex-specific reaction norms to intraspecific larval competition in the mosquito Aedes aegypti. J Evol Biol 16, 721–730 (2003).

33. Hochberg, M. E., Michalakis, Y. & De Meeus, T. Parasitism as a constraint on the rate of life-history evolution. J Evol Biol 5, 491–504 (1992).

34. Michalakis, Y. & Hochberg, M. E. Effets des parasites sur les traits biodémographiques de leurs hotes: Une revue des études récentes. Parasite 1, 291–294 (1994).

35. Agnew, P. & Koella, J. C. Life history interactions with environmental conditions in a host-parasite relationship and the parasite’s mode of transmission. Evol Ecol 13, 67–91 (1999).

36. Duneau, D. & Ebert, D. Host sexual dimorphism and parasite adaptation. PLoS Biol 10, (2012).

37. Vincent, C. M. & Sharp, N. P. Sexual antagonism for resistance and tolerance to infection in Drosophila melanogaster. Proceedings of the Royal Society B: Biological Sciences 281, (2014).

38. Klein, S. L. Hormonal and immunological mechanisms mediating sex differences in parasite infection. Parasite Immunology vol. 26 247–264 Preprint at 10.1111/j.0141-9838.2004.00710.x (2004).

39. Nunn, C. L., Lindenfors, P., Pursall, E. R. & Rolff, J. On sexual dimorphism in immune function. Philosophical Transactions of the Royal Society B: Biological Sciences 364, 61–69 (2009).

40. Restif, O. & Amos, W. The evolution of sex-specific immune defences. Proceedings of the Royal Society B: Biological Sciences 277, 2247–2255 (2010).

41. Bacelar, F. S., White, A. & Boots, M. Life history and mating systems select for male biased parasitism mediated through natural selection and ecological feedbacks. J Theor Biol 269, 131–137 (2011).

42. Barletta Ferreira, A. B. et al. Sexual Dimorphism in Immune Responses and Infection Resistance in Aedes aegypti and Other Hematophagous Insect Vectors. Frontiers in Tropical Diseases 3, (2022).

43. Schlenke, T. A., Morales, J., Govind, S. & Clark, A. G. Contrasting infection strategies in generalist and specialist wasp parasitoids of Drosophila melanogaster. PLoS Pathog 3, 1486–1501 (2007).

44. Dharmarajan, G., Gupta, P., Vishnudas, C. K. & Robin, V. V. Anthropogenic disturbance favours generalist over specialist parasites in bird communities: Implications for risk of disease emergence. Ecology Letters vol. 24 1859–1868 Preprint at 10.1111/ele.13818 (2021).

45. Sprague, V. Annotated List of Species of Microsporidia. in Comparative Pathobiology 31–334 (1977). 10.1007/978-1-4613-4205-2_2.

46. Solter, L. F. Epizootiology of Microsporidiosis in Invertebrate Hosts. Microsporidia: Pathogens of Opportunity 165–194 (2014).

47. Andreadis, T. G. Evolutionary strategies and adaptations for survival between mosquito-parasitic microsporidia and their intermediate copepod hosts: a comparative examination of Amblyospora connecticus and Hyalinocysta chapmani (Microsporidia: Amblyosporidae). Folia Parasitol (Praha) 52, 23 (2005).

48. Vulule, J. M. et al. Reduced susceptibility of Anopheles gambiae to permethrin associated with the use of permethrin-impregnated bednets and curtains in Kenya. Med Vet Entomol 8, 71–75 (1994).

49. Kelly, J. F., Anthony, D. W. & Dillard, C. R. A laboratory evaluation of the microsporidian Vavraia culicis as an agent for mosquito control. J Invertebr Pathol 37, 117–122 (1981).

50. Fukuda, T., Willis, O. R. & Barnard, D. R. Parasites of the Asian tiger mosquito and other container-inhabiting mosquitoes (Diptera: Culicidae) in northcentral Florida. J Med Entomol 34, 226–233 (1997).

51. Zeller, M. & Koella, J. C. Effects of food variability on growth and reproduction of A edes aegypti. Ecol Evol 6, 552–559 (2016).

52. Kulma, K., Saddler, A. & Koella, J. C. Effects of age and larval nutrition on phenotypic expression of insecticide-resistance in Anopheles mosquitoes. PLoS One 8, e58322 (2013).

53. Dao, A. et al. Reproduction-longevity trade-off in Anopheles gambiae (Diptera: Culicidae). J Med Entomol 47, 769–777 (2010).

54. Koella, J. C. & Lyimo, E. O. Variability in the relationship between weight and wing length of Anopheles gambiae (Diptera: Culicidae). J Med Entomol 33, 261–264 (1996).

55. Petersen, V. et al. Assessment of the correlation between wing size and body weight in captive Culex quinquefasciatus. Rev Soc Bras Med Trop 49, 508–511 (2016).

56. Abràmoff, M. D., Magalhães, P. J. & Ram, S. J. Image processing with ImageJ. Biophotonics international 11, 36–42 (2004).

57. Team, R. C. R: A language and environment for statistical com-puting. Preprint at (2017).

58. Hartig, F. DHARMa: Residual Diagnostics for Hierarchical (Multi-Level / Mixed) Regression Models. Preprint at https://CRAN.R-project.org/package=DHARMa (2022).

59. Fox, J. & Weisberg, S. Using car and effects Functions in Other Functions. Published online 1–5 (2020).

60. Bates, D., Mächler, M., Bolker, B. & Walker, S. Fitting Linear Mixed-Effects Models using lme4. (2014).

61. Lenth, R. V. emmeans: Estimated Marginal Means, aka Least-Squares Means. Preprint at https://CRAN.R-project.org/package=emmeans (2023).

62. Hothorn, T., Bretz, F. & Westfall, P. Simultaneous inference in general parametric models. Biom J 50, 346–363 (2008).

