## Supplementary information for "Host-specific effects of a generalist parasite of mosquitoes" by Tiago G. Zeferino and Jacob C. Koella for "Host-specific effects of a generalist parasite of mosquitoes"

**Table S1.** Larvae and pupae mortality.

**Table S2.** Age at pupation and wing length.

**Table S3.** Sex ratio.

**Table S4.** Presence of spores and spore density.

**Table S1. Larvae and pupae mortality.**

| Tested effect | | | |
|---|---|---|---|
| **Model 1a -** Larvae mortality | **df** | $\chi^2$ | ***p*** |
| Species | 1 | 523.82 | **<0.001** |
| Exposure | 1 | 112.25 | **<0.001** |
| Species : Exposure | 1 | 0.02 | 0.888 |
| **Model 1b -** Pupae mortality | **df** | $\chi^2$ | ***p*** |
| Species | 1 | 11.34 | **<0.001** |
| Exposure | 1 | 2.47 | 0.115 |
| Species : Exposure | 1 | 1.29 | 0.255 |

**Table S2. Age at pupation and wing length.**

| **Tested effect** | | | | |
|---|---|---|---|---|
| **Model 2a** – Age at pupation | **df** | **Mean Sq** | **F** | ***p*** |
| Species | 1 | 189.6 | 298.32 | **<0.001** |
| Exposure | 1 | 94.0 | 147.90 | **<0.001** |
| Sex | 1 | 37.4 | 58.88 | **<0.001** |
| Species : Exposure | 1 | 23.2 | 36.56 | **<0.001** |
| Species : Sex | 1 | 4.0 | 6.25 | **0.012** |
| Exposure : Sex | 1 | 2.7 | 4.27 | **0.038** |
| Species : Exposure : Sex | 1 | 1.2 | 1.90 | 0.168 |
| **Model 2b –** Wing length | **df** | **Mean Sq** | **F** | ***p*** |
| Species | 1 | 61.9 | 880.08 | **<0.001** |
| Exposure | 1 | 7.0 | 100.46 | **<0.001** |
| Sex | 1 | 31.5 | 448.23 | **<0.001** |
| Species : Exposure | 1 | 1.3 | 18.62 | **<0.001** |
| Species : Sex | 1 | 5.8 | 82.88 | **<0.001** |
| Exposure : Sex | 1 | 0.0 | 0.00 | 0.972 |
| Species : Exposure : Sex | 1 | 0.1 | 1.36 | 0.242 |

**Table S3. Sex ratio.**

| **Tested effect** | | | |
|---|---|---|---|
| **Model 3a** – Sex ratio | **df** | $\chi^2$ | ***p*** |
| Species | 1 | 14.61 | **<0.001** |
| Exposure | 1 | 28.63 | **<0.001** |
| Age at pupation | 1 | 97.00 | **<0.001** |
| Species : Exposure | 1 | 10.34 | **0.001** |
| Species : Age at pupation | 1 | 22.58 | **<0.001** |
| Exposure : Age at pupation | 1 | 33.47 | **<0.001** |
| Species : Exposure : Age at pupation | 1 | 15.10 | **<0.001** |

**Table S4. Presence of spores and spore density.**

| **Tested effect** | | | |
|---|---|---|---|
| **Model 4a** – Presence of spores (*Aedes*) | **df** | $\chi^2$ | ***p*** |
| Age | 1 | 3.99 | **0.046** |
| Sex | 1 | 1.28 | 0.258 |
| Age : Sex | 1 | 0.02 | 0.895 |
| **Model 4b –** Presence of spores (*Anopheles*) | | | |
| Age | 1 | 501.28 | **<0.001** |
| Sex | 1 | 0.25 | 0.618 |
| Age : Sex | 1 | 0.08 | 0.783 |
| **Model 4c** – Spore density (*Aedes*) | | | |
| Age | 1 | 19.68 | **<0.001** |
| Sex | 1 | 1.07 | 0.300 |
| Age : Sex | 1 | 6.07 | **0.014** |
| **Model 4d –** Spore density (*Anopheles*) | | | |
| Age | 1 | 165.84 | **<0.001** |
| Sex | 1 | 7.22 | **0.007** |
| Age : Sex | 1 | 3.44 | 0.064 |
